# Wedding higher taxonomic ranks with metabolic signatures coded in prokaryotic genomes

**DOI:** 10.1101/044115

**Authors:** Gregorio Iraola, Hugo Naya

## Abstract

Taxonomy of prokaryotes has remained a controversial discipline due to the extreme plasticity of microorganisms, causing inconsistencies between phenotypic and genotypic classifications. The genomics era has enhanced taxonomy but also opened new debates about the best practices for incorporating genomic data into polyphasic taxonomy protocols, which are fairly biased towards the identification of bacterial species. Here we use an extensive dataset of Archaea and Bacteria to prove that metabolic signatures coded in their genomes are informative traits that allow to accurately classify organisms coherently to higher taxonomic ranks, and to associate functional features with the definition of taxa. Our results support the ecological coherence of higher taxonomic ranks and reconciles taxonomy with traditional chemotaxonomic traits inferred from genomes. KARL, a simple and free tool useful for assisting polyphasic taxonomy or to perform functional prospections is also presented (https://github.com/giraola/KARL).

---

In 1735, Carl von Linné released the Systema Naturae ^1^ setting a cornerstone in biology by establishing a formal system for the unambiguous nomenclature of living things, underpinning the sciences of taxonomy and systematics. However, these disciplines were originally conceived for eukaryotic organisms where classification was mainly based on morphological macroscopic traits, suffering from expectable inconsistencies when analogous principles were applied to prokaryotes. This caused that the classification prokaryotes remained a controversial field restricted to a small number of experts highly trained to develop and reproduce tuned-up chemotaxonomic and phenotypic tests. Afterwards, the pioneering work of Carl Woese in the mid-1980s caused a sudden change which shed light onto the evolutionary history of prokaryotes by introducing the 16S gene analysis ^2^. Since then, many molecular characterization tools have been developed but the state-of-the-art strategy for assigning prokaryotes into novel or already described taxonomic units relies on polyphasic taxonomy, an approach which tries to integrate all available chemotaxonomic and genotypic information to build a classification consensus ^3^.

The advent of high-throughput sequencing (HTS) has allowed the incorporation of genomic information into polyphasic taxonomy ^4^, but much effort has been invested to define classification rules for species in detriment of higher taxonomic ranks. Recently, two approaches that overcome this problem have been published: i) PhyloPhlAn constructs a high-resolution phylogenetic tree using a previously optimized set of 400 marker genes that accurately defines most taxa ^5^ and, ii) Microtaxi identifies taxon-specific genes and relies on a simple counting scheme to assign genomes to each taxon ^6^. Both approaches were optimized with a subset of the available genomic diversity (around 2, 000), do not provide automatic updating alternatives as new genomes become available and, fundamentally, do not allow to associate the taxonomic position with distinctive functional features of organisms.

In the present work we submit the hypothesis that higher taxonomic ranks (domain to genus) can be inferred from analyzing metabolic signatures of genomes. In turn, this hypothesis arises from the notion that higher ranks are ecologically coherent, meaning that most organisms within the same hierarchical level should display certain rationality in their lifestyles and ecological traits ^7^. This is supported by the underlying signal of vertical evolution found in genes coding basal functions used as taxonomic markers ^5, 8^. Indeed, signature metabolic genes that define taxon-specific ecological traits should be recognizable and used to define taxonomic ranks, similar to the approach implemented in Microtaxi ^6^, but also considering that the ecological coherence of high taxa could result from unique gene combinations, without any of them being taxon specific ^7^. This requires the implementation of more powerful methods to capture informative patterns in highly-dimentional spaces.

To prove this ecological coherence of taxa at higher ranks and the usefulness of metabolic features as taxonomic markers, we used an extensive dataset of 33, 236 archaeal and bacterial genomes representing 2 domains, 55 phyla, 67 classes, 163 orders, 328 families and 1,480 genera. For each genome the presence orabsence of 1, 328 different enzymes was assessed by parsing their annotations. Fig. 1a exemplifies how principal components analysis (PCA) resulting from enzyme patterns spatially discriminate taxa at different ranks. This separation is coherent with highly frequent or infrequent enzymes (Fig. 1b) and with Jaccard distance-based clustering (Fig. 1c). Then, we take the families Helicobacteraceae (intestinal gram-negatives associated with Crohn’s disease ^9^) and Enterococcaceae (intestinal gram-positives associated to probiotic effects^10^) to illustrate that enzyme patterns can cluster genomes according to their taxonomic position (Fig. 2a) based on the presence of distinctive combinations and subsets of marker enzymes (Fig. 2b). Beyond that identifying these single markers can provide useful information about taxon-specific molecular functions, we show that this information is also scalable to metabolic pathways allowing the isolation of those that significantly distinguish them (Fig. 2c). In Fig. 2d we show the full reference metabolism for starch and sucrose, which is one out of eight pathways that significantly distinguish both families each other (Fig. 2c), evidencing that the members of Enterococcaceae family present a vast distribution of enzymes while it is much more limited for the Helicobacteraceae genomes. This kind of functional prospection uncovers a strong link between metabolic potential and taxonomy, which indeed has been evidenced recently by modeling the variation of metabolomic data and community composition using metagenomic data ^11^.

**Figure 1.**
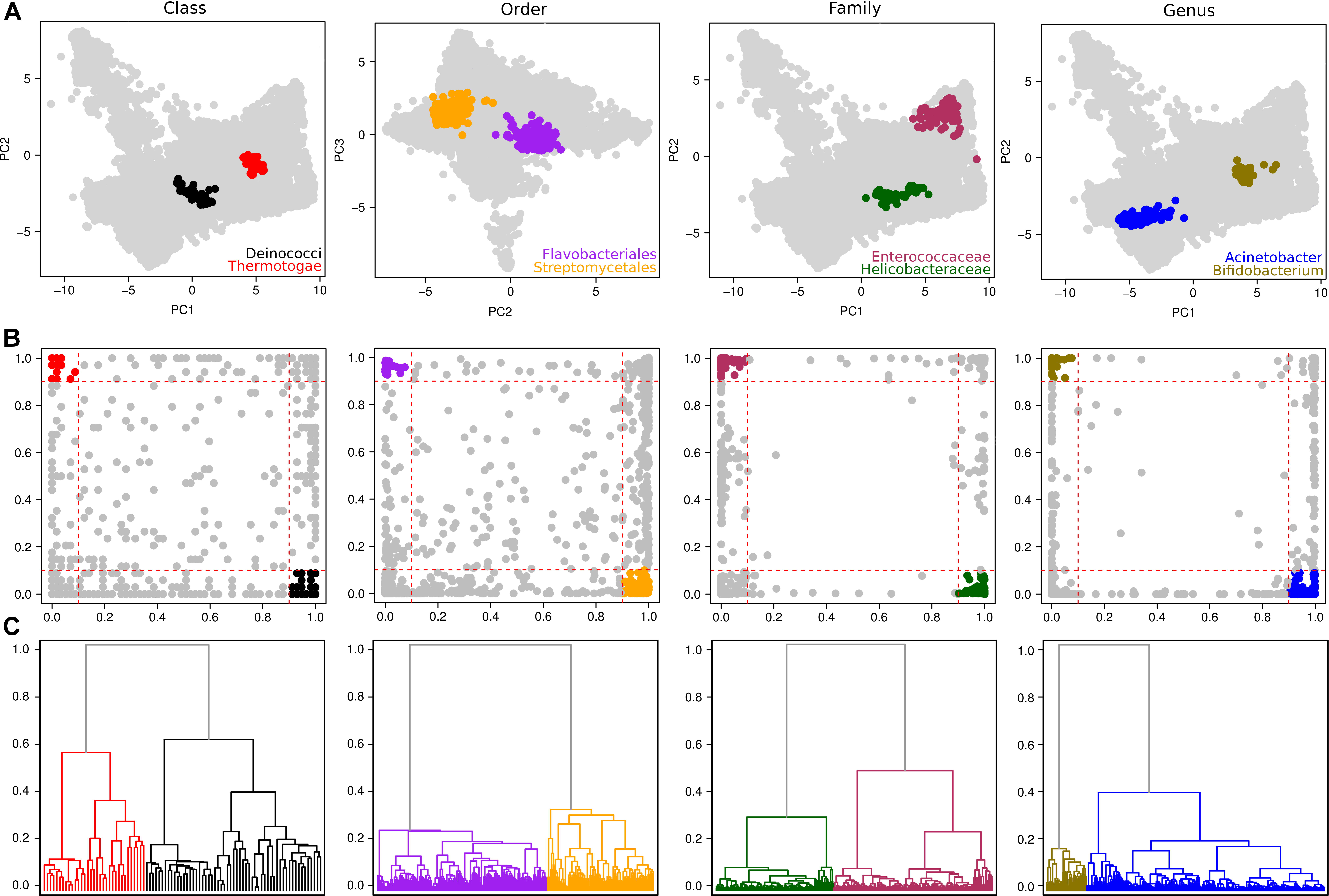
Informativeness of enzyme patterns. A) Principal Component Analysis using enzyme presence/absence vectors shows the discrimination of different pairs of taxa at every rank. B) For the same pairs of taxa, the frequency of each enzyme is plotted. Enzymes with frequency >0.9 in one taxon and <0.1 in the other and viceversa are highlighted. C) Cladogram based on pairwise Jaccard’s distances.

**Figure 2.**
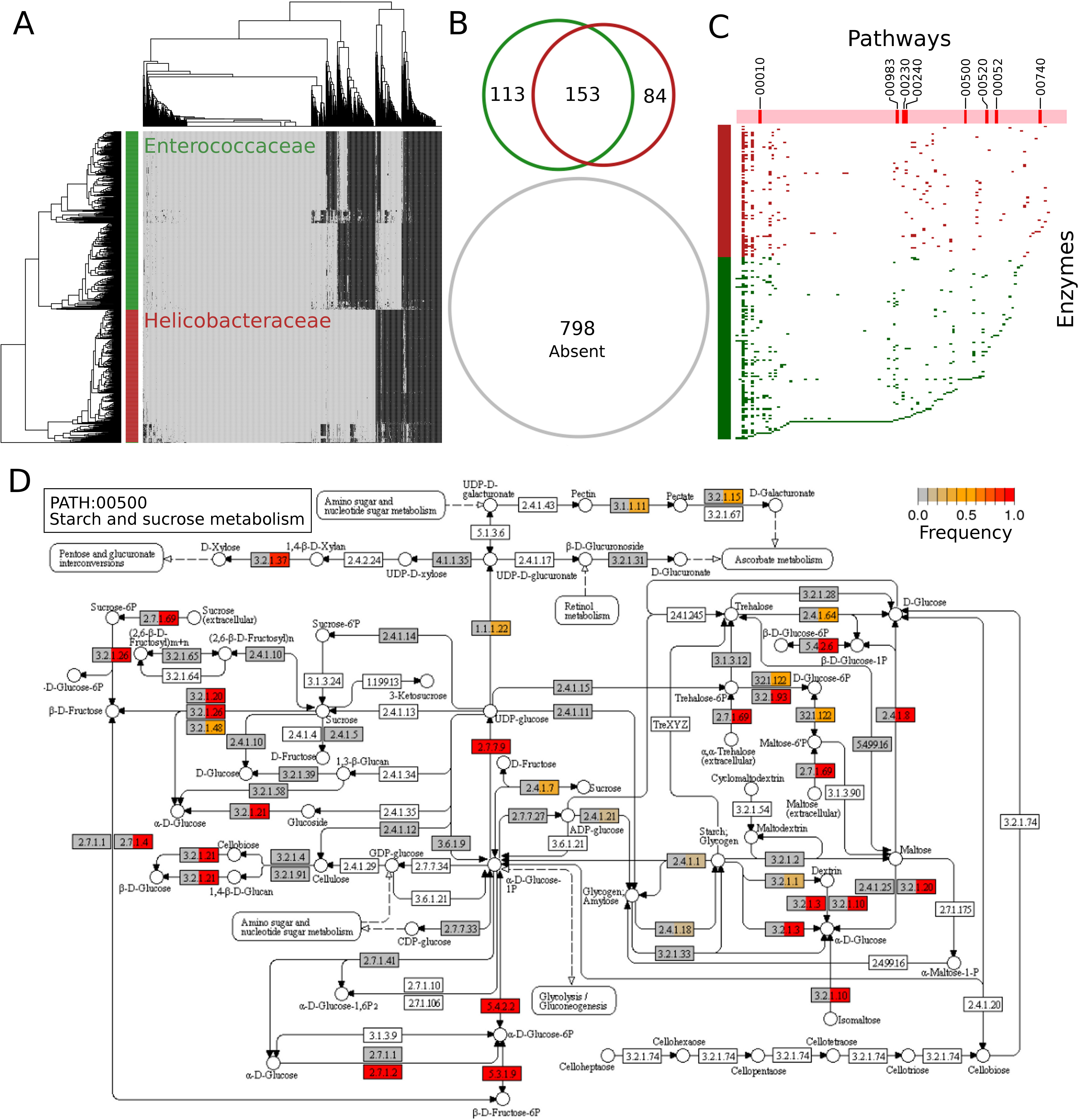
Example on families Helicobacteraceae and Enterococcacae. A) Heatmap showing the hierarchical clustering of genomes belonging to both families, exhibiting taxon-specific enzyme clusters. B) Venn diagram showing the distribution of enzymes between families. C) Distribution of enzymes inside metabolic pathways. Pathways that significantly differ (see Supplementary Methods) between families are labeled according to KEGG pathway identifiers. D) KEGG reference pathway for starch and sucrose metabolism. Each enzyme box is divided in two: left side for Helicobacteraceae and right side for Enterococcaceae. The frequency of each enzyme in each taxon is colored from grey (0) to red (1).

As we showed that enzyme patterns are enough informative to discriminate taxa at different ranks and given the binary nature of this data, we built support vector machine (SVM) classification models using linear kernels by splitting the data in subsets corresponding to each taxa against the rest. All models were 10-fold corss-validated and performed very well independently of the taxonomic rank, reaching median precisions above 90%, false positive rates below 2% and false negative rates below 5% (Supplementary Fig. 1). The very low rate of false positives is important since the practical cost of assigning a strain into a certain taxon is higher than keeping it as unknown. The explanation for non-optimal performance is the biased number of available genomes per taxa, since strong correlations (R^2^ = 0.76, p-value = 2 × 10^−16^) were found between the number of genomes (at every rank) and classification performance measured as previously (Supplementary Fig. 2). Additionally, when considering taxa with increasing minimal number of genomes all performance parameters rapidly scale to optimal values. For example, when all taxa at family rank are included, the models showed a median precision of 94% and the interquartile range (IQR) between 72% and 100%, however when looking at models with at least 10 genomes the first quartile increased to 89% and the median to 95%, and for models with more than 50 genomes the first quartile scaled to 94% and the median to 97%. This tendency was observed for all ranks (Supplementary Fig. 3) and demonstrated that classification errors are better explained due to biased data than to lack of information in enzyme patterns. Indeed, for some small taxa like classes Archaeoglobi (n = 7), Chlorobia (n = 12), Thermodesulfobacteria (n = 8) or Dictyoglomia (n = 2) classification precision was 100%. Interestingly, most of these taxa exhibit powerful ecological constraints reflected in very informative enzyme patterns that supports the ecological coherence of higher ranks and its association to genome-encoded signatures, overcoming their low representation in the whole dataset. Anyway, these observations reinforce the importance of sequencing genomes not only for the anthropocentric convenience but also for mere taxonomic interest, and also warrant the optimization of our approach as new genomes become available.

**Figure 3.**
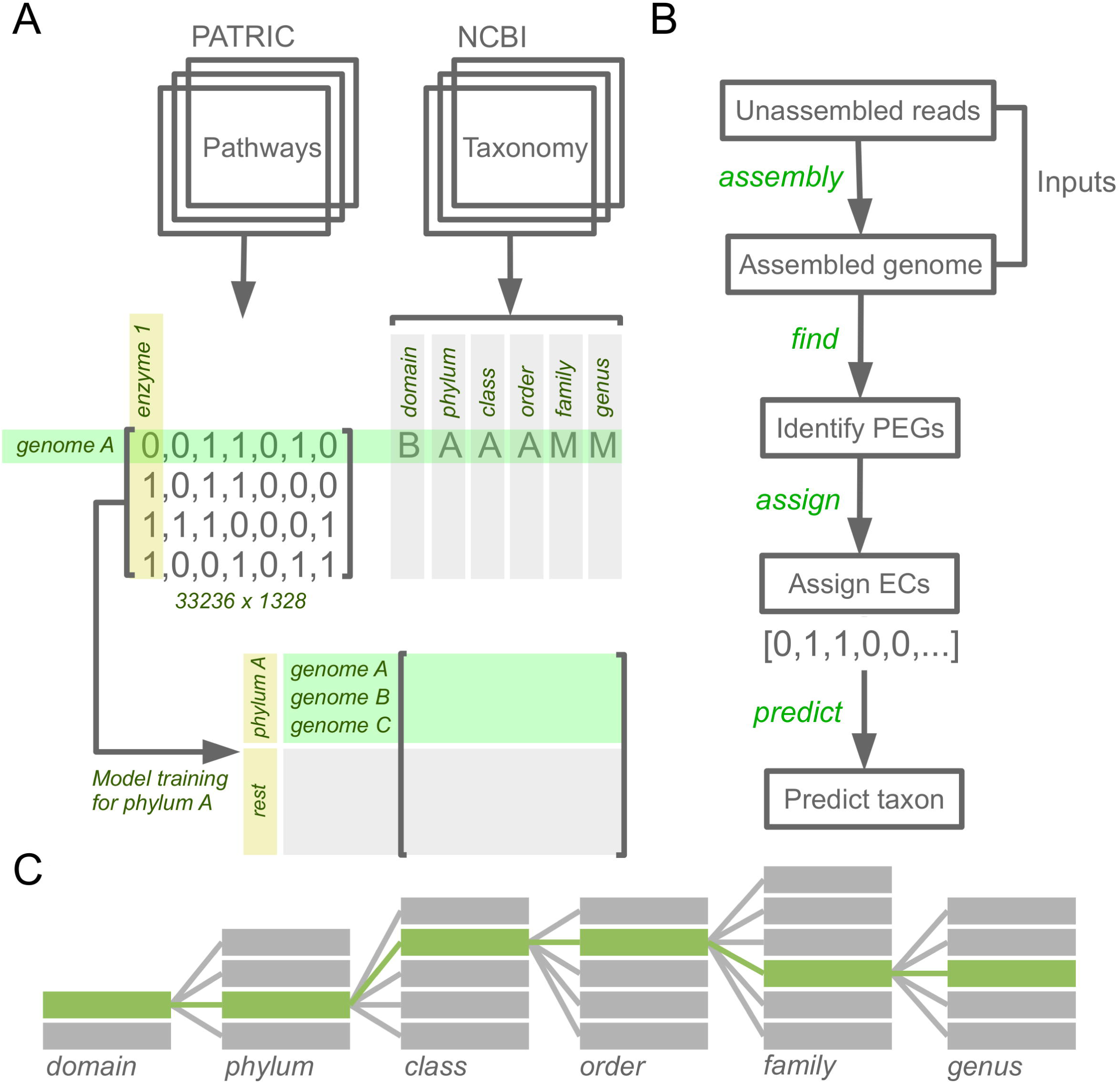
KARL pipeline. A) The workflow for building the dataset from genomes and annotations. B) Step-by-step analysis to classify a new genome. C) Schematic representation of included taxonomic ranks.

The robustness of predictions was further tested by applying the algorithm to an external set of 108 genomes obtained from very recent issues of Genome Announcements (http://genomea.asm.org/), hence not used in any step of model construction. At genus rank the average precision was 92% (Supplementary Tab. 1), holding that obtained with cross-validated models. Interestingly, all misclassified genomes (n = 8) were predicted as unknown instead as any known genus erroneously, reinforcing the resilience of classification models against false positives. Additionally, the classification was totally consistent for genomes whose genus was truly unknown. For example, the bacilli bacterium VTI3104 was assigned within an unknown genus inside the Bacillaceae family, in accordance to its divergent phylogenetic position inside bacilli ^12^. Alike, the Oscillatoriales cyanobacterium MTP1 was in fact classified within the Oscillatoriales order but was assigned to an unknown family and genus, in accordance to the low 16S identity (~90%) against its closest neighbors ^13^. At higher ranks, classification was perfect for those genomes which were correctly assigned at genus rank, since the algorithm takes benefit from the hierarchical structure of taxonomy and stops once the sample has been assigned to a certain genus by completing the lineage with precomputed information. Alike, when these genomes were tested for selected ranks above genus the results were totally coherent with the corresponding taxon. Seven out of 8 genomes that were incorrectly assigned at genus rank were then correctly classified at family rank. No misclassifications were observed at order, class, phylum and domain ranks.

To ease the straightforward use of all models and datasets developed here, we built an R ^14^ package called KARL (https://github.com/giraola/KARL) that allows the user to predict the membership of any newly sequenced genome to each high taxonomic rank by inputing just HTS reads, assembled genomes or annotation files. Additionally, the user can explore and improve each taxon-specific classification model by performing automatic feature selection procedures and compare between same rank taxa. A handful of functions allows the identification, comparison and visualization of meaningful metabolic signatures among taxa, including the automated production of all graphs and illustrations herein presented. Finally, it can connect the PATRIC database ^15^ to automatically update models and datasets based on newly released genomes. The full KARL pipeline is shown in Fig. 3. A detailed user guide explains theoretical and practical aspects from installation to step-by-step implementation of all features herein described (Supplementary Methods). KARL is intended to be a useful tool for assisting polyphasic taxonomy and to perform functional prospections and comparisons of prokaryotic genomes.

## Methods

### Genomic data

Annotation files were accessed from the PATRIC database ^15^ for a total of 33, 236 available prokaryotic genomes. In-house R ^14^ scripts were designed to extract enzymes and pathways datafrom eachfileand build a presence/absence matrix; each columninthe matrix represents one out of 1,328 different enzymes detected in at least one genome and identified with its unique EC number. The taxize R package ^16^ was used to retrieve the domain, phylum, class, order, family and genus for each genome from the NCBI Taxonomy database.

### Classification models

RWeka ^17^ was used to train Support Vector Machine (SVM) models by splitting the data in two categories: one containing the considered taxon and other with the rest. For all cases, given the binary nature of data, a linear kernel function was preferred ^18^. Each model can be improved by performing a feature selection scheme that involved three steps: i) attributes (enzymes) with a frequency lower than 0.1 or greater than 0.9 in both categories were removed, ii) over the remaining set highly correlated attributes (>0.9) were removed just keeping those with lower average correlation values and, iii) Information Gain ratios were calculated for each attribute and a Davies test for slope change was applied to identify the Gain Ratio cut-off for keeping those most informative enzymes (Supplementary Methods). Evaluation was initially performed by implementing a 3-repeated 10-fold cross-validation scheme to each model and then misclassified genomes were evaluated using a metric based on the integration of Hamming distances in the PCA space (Supplemental Methods). Further testing was performed with an external dataset (Supplementary Tab. 1).

### Taxonomy prediction

The first step for predicting the membership of a new genome into any taxonomic unit implies to determine its presence/absence pattern for the 1, 328 enzymes tested. Assessing this depends on the input: for unassembled sequencing reads the SPAdes genome assembler ^19^ generates a *de novo* assembly, else, if the input is a draft or finished genome the pipeline starts using Prodigal ^20^ and/or Glimmer ^21^ to predict protein-encoding genes. Finally, BLASTp ^22^ identifies enzymes presence/absence by searching against individual databases built from FIGfams ^23^ and KEGG Orthology ^24^. A certain enzyme is considered present if there is any BLASTp hit with identity >70%, query coverage >90% and e-value <0.001 (these values were selected based on a grid search analysis that evaluated the classification performance using combinations of identity from 25% to 95% and query coverage from 50% to 95%) (Supplementary Fig. 4). The presence/absence vector is then inputed to each taxon-specific SVM model.

### KARL package

The whole methods were implemented as an R package called KARL freely available at GitHub repository (https://github.com/giraola/KARL). This package allows the standalone implementation of the whole applications described here in three operational modules: Explorer, Predictor and Updater. Explorer implements a handful of functions for automatic comparison of taxa, allowing to identify metabolic signatures at enzyme and pathway levels. Predictor allows to predict taxonomy from sequencing reads, assembled or annotated genomes, evaluate predictions and optimize classification models. Updater allows to automatically update datasets and models with new available sequenced genomes in public databases. An in-depth description of practical and theoretical aspects are provided in the full user manual (Supplemental Methods).

## Acknowledgements

This work has been partially funded by Fondo de Convergencia Estructural del Mercosur (FOCEM) grant COF 03/11, Comisión Sectorial de Investigación Cientf́ica (CSIC) and Agencia Nacional de Investigatión e Innovatión (ANII) grant FSSA_X_2014_1_105252.

## Competing Interests

The authors declare that they have no competing financial interests.

## Correspondence

Correspondence and requests for materials should be addressed to H.N. or to G.I.

## References

1. Linneaus, C. v. Systema naturae., vol. 1 (Holmiae:Impensis Direct. Laurentii Salvii, 1758).

2. Woese, C. R., Kandler, O. & Wheelis, M. L. Towards a natural system of organisms: proposal for the domains Archaea, Bacteria, and Eucarya. Proc. Natl. Acad. Sci. U.S.A. 87, 4576–4579 (1990).

3. Vandamme, P. et al. Polyphasic taxonomy, a consensus approach to bacterial systematics. Microbiol. Rev. 60, 407–438 (1996).

4. Kim, M., Oh, H. S., Park, S. C. & Chun, J. Towards a taxonomic coherence between average nucleotide identity and 16S rRNA gene sequence similarity for species demarcation of prokaryotes. Int. J. Syst. Evol. Microbiol. 64, 346–351 (2014).

5. Segata, N., Bornigen, D., Morgan, X. C. & Huttenhower, C. PhyloPhlAn is a new method for improved phylogenetic and taxonomic placement of microbes. Nat Commun 4, 2304 (2013).

6. Gupta, A. & Sharma, V. K. Using the taxon-specific genes for the taxonomic classification of bacterial genomes. BMC Genomics 16, 396 (2015).

7. Philippot, L. et al. The ecological coherence of high bacterial taxonomic ranks. Nat. Rev. Microbiol. 8, 523–529 (2010).

8. Snel, B., Bork, P. & Huynen, M. A. Genome phylogeny based on gene content. Nat. Genet. 21, 108–110(1999).

9. Kaakoush, N. O. et al. Detection of Helicobacteraceae in intestinal biopsies of children with Crohn’s disease. Helicobacter 15, 549–557 (2010).

10. Lory, S. The Family Enterococcaceae, vol. 1 (Springer, 2014).

11. Noecker, C. et al. Metabolic model-based integration of microbiome taxonomic and metabolomic profiles elucidates mechanistic links between ecological and metabolic variation. mSystems 1 (2016). URL http://msystems.asm.org/content/1/1/e00013-15. http://msystems.asm.org/content/1/1/e00013-15.full.pdf.

12. Tetz, G. & Tetz, V. Complete Genome Sequence of Bacilli bacterium Strain VT-13-104 Isolated from the Intestine of a Patient with Duodenal Cancer. Genome Announc 3 (2015).

13. Hallenbeck, P. C., Grogger, M., Mraz, M. & Veverka, D. Draft Genome Sequence of a Thermophilic Cyanobacterium from the Family Oscillatoriales (Strain MTP1) from the Chalk River, Colorado. Genome Announc 4 (2016).

14. R Development Core Team. R: A Language and Environment for Statistical Computing. R Foundation for Statistical Computing, Vienna, Austria (2008). URL http://www.R-project.org. ISBN 3-900051-07-0.

15. Wattam, A. R. et al. PATRIC, the bacterial bioinformatics database and analysis resource. Nucleic Acids Res. 42, D581–591 (2014).

16. Chamberlain, S. A. & Szocs, E. taxize: taxonomic search and retrieval in R. F1000Res 2, 191 (2013).

17. Hornik, K., Zeileis, A., Hothorn, T. & Buchta, C. Rweka: an r interface to weka. R package version 0.3-2 (2007).

18. Iraola, G., Vazquez, G., Spangenberg, L. & Naya, H. Reduced set of virulence genes allows high accuracy prediction of bacterial pathogenicity in humans. PLoS ONE 7, e42144 (2012).

19. Bankevich, A. et al. SPAdes: a new genome assembly algorithm and its applications to singlecell sequencing. J. Comput. Biol. 19, 455–477 (2012).

20. Hyatt, D. et al. Prodigal: prokaryotic gene recognition and translation initiation site identification. BMC Bioinformatics 11, 119 (2010).

21. Delcher, A. L., Harmon, D., Kasif, S., White, O. & Salzberg, S. L. Improved microbial gene identification with GLIMMER. Nucleic Acids Res. 27, 4636–4641 (1999).

22. Altschul, S. F., Gish, W., Miller, W., Myers, E. W. & Lipman, D. J. Basic local alignment search tool. J. Mol. Biol. 215, 403–410 (1990).

23. Meyer, F., Overbeek, R. & Rodriguez, A. FIGfams: yet another set of protein families. Nucleic Acids Res. 37, 6643–6654 (2009).

24. Tanabe, M. & Kanehisa, M. Using the KEGG database resource. Curr Protoc Bioinformatics Chapter 1, Unit1.12 (2012).

